# Development of Equine Immunoglobulin Fragment F(ab’)_2_ with High Neutralizing Capability against SARS-CoV-2

**DOI:** 10.1101/2021.03.09.434030

**Authors:** Divya Gupta, Farhan Ahmed, Dixit Tandel, Haripriya Parthasarathy, Dhiviya Vedagiri, Vishal Sah, B Krishna Mohan, Siddarth Daga, Rafiq Ahmad Khan, Chiranjeevi Kondiparthi, Prabhudas Savari, Sandesh Jain, Jaya Daga, Shashikala Reddy, Nooruddin Khan, Krishnan Harinivas Harshan

## Abstract

The ongoing pandemic, COVID-19, caused by SARS-CoV-2 has taken the world, and especially the scientific community by storm. While vaccines are being introduced into the market, there is also a pressing need to find potential drugs and therapeutic modules. Remdesivir is one of the antivirals currently being used with a limited window of action. As more drugs are being vetted, passive immunotherapy in the form of neutralizing antibodies can provide immediate action to combat the increasing numbers of COVID-positive cases. Herein, we demonstrate that equines hyper-immunized with chemically inactivated SARS-CoV-2 generate high titers of antibody with a strong virus neutralizing potential. ELISA performed with pooled antisera displayed highest immunoglobulin titer on 42 days post-immunization, at 1:51,200 dilutions. F(ab’)_2_ immunoglobulin fragments generated from the pools also showed very high, antigen-specific affinity at 1:102,400 dilutions. Finally, *in vitro* virus neutralization assays confirmed that different pools of F(ab’)_2_ fragments could successfully neutralize SARS-CoV-2 with titers well above 25,000, indicating the potential of this strategy in treating severe COVID-19 cases with high titers. The F(ab’)_2_ was able to cross neutralize another SARS-CoV-2 strain, demonstrating its efficacy against the emerging viral variants and the importance of this approach in our efforts of eradication of COVID-19. In conclusion, this study demonstrates that virus-neutralizing antibodies raised in equines can potentially be used as a treatment regimen in the form of effective passive immunotherapy to combat COVID-19.

## INTRODUCTION

The ongoing pandemic COVID-19 caused by severe acute respiratory syndrome coronavirus-2 (SARS-CoV-2) has caused over 115 million total infections with more than 2.5 million deaths globally to date^1^. The severity and the scale of the pandemic have imposed an unprecedented strain on human health and the global economy. Though several vaccines have been approved for immunization, it would require years of continuous vaccination drive before we defeat the disease^2,3^. Remdesivir is an antiviral drug used for treating COVID-19, though with limited efficacy which can only shorten the period of hospitalization if administered at the early phase of infection^4^. The long delay in vaccination programs coupled with the absence of effective drug indicates that COVD-19 is far from being over^5–7^. The situation clearly calls for multiple approaches in countering the viral spread.

Neutralizing antibody (nAb)-based passive immunotherapy has been used as an antiviral therapy module against various intractable viral diseases^8^ by blocking the viral attachment and entry into the host cell. In the pandemic setting, convalescent plasma from the recovered patient has been used as an emergency treatment plan for the emerging virus infectious diseases^9^ and congruously, it has been approved by USFDA in the COVID-19 too^10^. While convalescent plasma is considered as a quick source of polyclonal nAb against the infectious agent, its scope is limited due to the lack of abundant blood source, heterogeneous antibody titer, and possible risks of transmission of blood-borne infections^7,11^. An alternative to convalescent plasma can be the antisera with improved efficacy obtained from hyper-immunized large animals such as equines as demonstrated against various infectious diseases and venoms^12–14^. Even though equine-derived antisera offer great potential for passive immunotherapy, they carry the risk of antibody-dependent enhancement of infection (ADE) and serum sickness^15,16^. To overcome this limitation, the next-generation passive immunotherapy uses the F(ab’)_2_ fragment of antigen-specific immunoglobulins, thus avoiding the risk of ADE by removing the Fc region of the antibody^17–19^.

Based on the above background, we have developed equine SARS-CoV-2 specific immunoglobulin fragment F(ab’)_2_ and evaluated its virus neutralization potential. In this process, the SARS-CoV-2 cultures of Indian isolate were propagated and chemically inactivated before immunizing equines for the evaluation of its immunogenicity and potency. The immunoglobulin fragments F(ab’)_2_ were prepared from the hyper-immunized equines and their virus neutralization potential was tested by microneutralization assay. The result of the study indicates the efficacy of SARS-CoV-2 specific F(ab’)_2_ fragments in the neutralization of the virus. This strategy is reproducible, easily scalable and could be made available for the masses. This approach of immunotherapy will considerably help in managing the global COVID-19 pandemic scenario.

## MATERIALS AND METHODS

### Cell Culture

Vero (CCL-81) cells were cultured in Dulbecco’s modified eagle medium (DMEM, Gibco) supplemented with 10% Fetal Bovine Serum (Hyclone) and 1 × Penicillin-Streptomycin cocktail (Gibco). Cells were continuously passaged at 70-80% confluency and were maintained in a humidified cell culture incubator at 37°C and 5% CO_2_.

### SARS-CoV-2 propagation, quantification, and infection

SARS-CoV-2 virus was isolated from COVID-19 patient sample. Briefly, viral transport media (VTM) with lower Ct values (<20) for SARS-CoV-2 Envelop and RdRp genes by real-time qRT-PCR were identified for culturing. 100μL of the filter-sterilized VTM was added to Vero cell monolayer in 96 well plates. Fresh media was supplemented three hrs post-infection (hpi) and the wells were observed daily for cytopathic effect (CPE). After the appearance of CPE, the supernatants of the corresponding wells were transferred to fresh monolayers of Vero cells and the culture was continued. The supernatants were regularly monitored by real-time qRT-PCR for viral genes as an indicator for viral replication. Two such isolates that established continuous replication were identified and sequenced by next-generation sequencing. Their genomic sequences have been deposited to the GISAID database ^20,21^ (GISAID ID: EPI_ISL_458075; virus ID-hCoV-19/India/TG-CCMB-O2-P1/2020, and EPI_ISL_458046; virus ID-hCoV-19/India/TG-CCMB-L1021/2020). These viral stocks were used for all the experiments in this study.

All viral cultures were maintained in serum-free media. Three days post-infection, the cell culture supernatant was collected, centrifuged at 5000 rpm for 10 min at 4°C to remove all the cell debris, and was stored at −80°C till further use. Infectious viral titers of the supernatants were measured by plaque forming assay (PFU/mL). All infections for the experimental assay were performed in serum-free media for two hrs at 37°C with the required amount of virus calculated from the respective PFU values.

### Real-time Quantification and Plaque forming assay

RNA was isolated using viral RNA isolation kit (MACHEREY-NAGEL GmbH & Co. KG). Real-time quantitative RT-PCR was performed in Roche LightCycler 480 either using a commercial kit (LabGun™ COVID-19 RT-PCR Kit; CV9032B) or following WHO guidelines using SuperScript™ III Platinum™ One-Step qRT-PCR Kit (Thermo Fisher) and TaqMan probes against SARS-CoV-2 E- and RdRp (Eurofins Scientific). Raw Ct values generated post analysis of qRT-PCR was used to score the supernatants. For plaque assay, 0.35 million Vero cells were seeded in 6 well plates and serial dilutions of virus supernatants (10^−3^ to 10^−8^) were used for infection in serum-free media. Two hrs post-infection, cells were briefly washed with 1 × PBS to remove unbound virus and overlaid with 1 × agarose overlay media (2 × DMEM, 5% FBS, 1% penicillin-streptomycin, 2% LMA). Plates were left undisturbed at 37°C with 5% CO_2_ in an incubator for 6-7 days. Later, 4% formaldehyde in 1 × PBS was added onto the overlay media for fixation and incubated for 15-20 min at 37°C. The overlay media along with formaldehyde were removed, the cells were washed briefly with 1× PBS and then stained with crystal violet stain (1% crystal violet in ethanol was used as the stock solution and 0.1% working solution was prepared in double distilled water). The plates were washed, dried and the number of clear zones in the plate was counted to determine the infectious titer as PFU/mL.

### Virus Inactivation

The cell culture supernatant containing SARS-CoV-2 was inactivated using beta propiolactone (BPL; HiMedia) at a ratio of 1:250 or 1:1000. After adding BPL to virus supernatants, the mixture was incubated at 4°C for 16 hrs with constant stirring, followed by four-hour incubation at 37°C to hydrolyze the remaining BPL in the solution. The inactivation of the virus was measured by plaque assay or CPE for three consecutive rounds. The absence of plaques or CPE in the lowest dilution confirmed the total inactivation. The BPL treated supernatants were precipitated by ultracentrifugation and the antigenic integrity of the samples was confirmed by immunoblotting.

### Immunoblotting

Infected cell lysates and virus supernatants were separated on SDS-PAGE gels to confirm the presence and integrity of viral proteins. All samples were lysed in a mild lysis buffer (Tris-Cl, NaCl, NP40; protease and phosphatase inhibitors) and Laemmli loading dye was added. Once the proteins were separated on the gels, they were transferred onto PVDF membranes for 2 hours and subsequently blocked in 5% BSA. Blots were probed with Nucleocapsid (1:8000) and Spike (1:2000) primary antibodies, and HRP-conjugated secondary antibodies. Image processing was performed using ImageJ^22^.

### Equine immunization plan

Separate groups (lots) each comprising of equines were formed and each lot was assigned a unique lot number. These unique numbers were used across the entire study involving activities such as immunization, bleeding and plasmapheresis. The immunization schedule comprised of primary immunization, during which period the animals were sensitized with inactivated SARS-CoV-2 viral antigen mixed with Freund’s complete adjuvant (FCA) as adjuvant and administered for a single time. The subsequent booster immunizations were administered with Freund’s incomplete adjuvant (FIA).

During the primary immunization, the animals were immunized on days 0, 15, 29, 35 and 45. On day zero, the equines were immunized with 1mL of viral antigen suspension containing 1 × 10^7^ inactivated virus particles mixed with equal volumes of FCA. The subsequent booster doses were prepared by mixing 0.5 mL of viral antigen (containing 1 × 10^7^ inactivated virus particles) with equal volume of FIA and administered periodically for boosting the immune response. Plasma samples from the immunized animals were tested periodically to estimate the antibody response against SARS-CoV-2 inactivated viral antigen. The highest dilutions of the plasma samples necessary to bind with specificity to viral antigen coated in the micro-titer plate were estimated by ELISA. Once the antibody response in the animals was saturated, they were bled and blood volumes equivalent to 1.5% of the individual body weight were collected in glass containers containing acid citrate dextrose solution (final concentration of 15% in the blood volume) to prevent the coagulation. The supernatant plasma in each of the containers was carefully collected and pooled for further studies.

### Measurement of serum IgG level and their titer

Antigen-specific total IgG was measured by indirect ELISA method. The whole viral antigen was coated in the 96 well plate (Nunc) using bicarbonate coating buffer (pH=9.5) overnight at 4°C. The coated plates were washed with washing buffer (0.05% Tween-20 in 1× PBS) and blocked with 4% skimmed milk solution for two hrs at RT followed by three rounds of washing. Sera from control and test group were added in the plate at 1:100 dilutions and incubated for two hrs at RT. Subsequently, the plate was washed four times and incubated with HRP conjugated anti-horse whole IgG secondary antibody (Sigma) for 1 hr at RT. After washing the plate five times, TMB substrate was added, the reaction continued for about three minutes, and stopped by addition of 0.2N H_2_SO_4_. Absorption maxima were recorded at 450nm and plotted on the XY axis graph.

In another set of experiment, IgG titer kinetics from 0 to 54 days post immunization were calculated by similar ELISA method with slight modification. Here, sera from ten animals for each time point were pooled in and serially diluted beginning from 1:100 to 1:204800 and added into viral antigen coated ELISA plates while rest of the steps were the same as above. Similar protocols were followed for the titration of F(ab’)_2_ fragments. Antibody titers were calculated by the reciprocal value of highest dilution at which absorbance value is ≥ twice the value of negative control in the same dilution series based on the earlier report^23^.

### Virus Neutralization assay

Neutralization capacities of the antisera and F(ab’)_2_ were measured by microneutralization assay in 96-well plates. For the initial standardization of the optimal number of viruses required for CPE in 100% wells, cells were infected in a 96 well format with the varying numbers of the virus particles. In all our neutralization studies, 300 virus particles were used for infection. For neutralization of virus by equine antisera, 30000 cells were seeded in each well of a 96 well plate 12 hrs before assay set up. 25 μL of serum-free media containing 300 infectious SARS-CoV-2 particles were mixed with 25μL of antiserum: serum-free media mix prepared separately. This mix contained antisera in 1:2, 1:4, 1:8 and up to 1:4096 parts of concentrations. The antisera: virus mixes were pre-incubated at 37°C for 1 hr before infection. Subsequently, the wells containing cells were washed with 1× PBS and the mixes were added to the corresponding wells. After the initial adsorption for 2 hrs at 37°C and 5% CO_2_, the virus containing media was replaced with fresh serum-sufficient media and incubated for six days. CPE developed in the wells were noted, media removed, and the remaining cells were fixed with 100μL of 4% formaldehyde at 37° C for 20 minutes. Post-fixation, formaldehyde was removed, wells were washed and the cells were stained with 0.1% trypan blue for 5 minutes to detect the live cells. The wells were observed against white light and scored for the presence or absence of CPE and CCID 50 was calculated by a modified Reed and Muench formula. The proportionate distance (PD) was first calculated using the formula (% positive above 50%-50%)/(% positive above 50%- % positive below 50%). The PD obtained was multiplied by the dilution below 50% and value obtained was added to the dilution below 50% to reach the dilution of CCID50).

### Preparation of F(ab’)_2_ immunoglobulin

Thirty liters of pooled plasma was subjected to enzymatic hydrolysis of IgG using pepsin (2% w/v) for 2 hrs with the pH adjusted to 3.3. The enzymatically treated plasma was subjected to complement inactivation by holding at a temperature of 56^°^C for 30 minutes. Further, caprylic acid was added gradually to make a final concentration of 5% v/v and mixed for one hour. Caprylic acid precipitates non-IgG proteins keeping the F(ab’)_2_ in solution. The antibody fragment F(ab’)_2_ in the supernatant was diafiltered and concentrated by ultra filtration through a 30 kDa cut-off membrane using 20 mM sodium acetate buffer with 0.9% sodium chloride. The resultant purified concentrated bulk becomes the key intermediate and tested for *in vitro* potency by ELISA and viral neutralization by the cell culture method. The concentrated bulk was formulated and filled as a final injectable dosage form, keeping the fill volume to 3 mL per vial. The finished product is intended for administration through either intramuscular or intravenous route based on the severity of the viral load and the urgency of the intervention. Immunization schedule along with the workflow is given in Figure 1.

**Fig.1.**
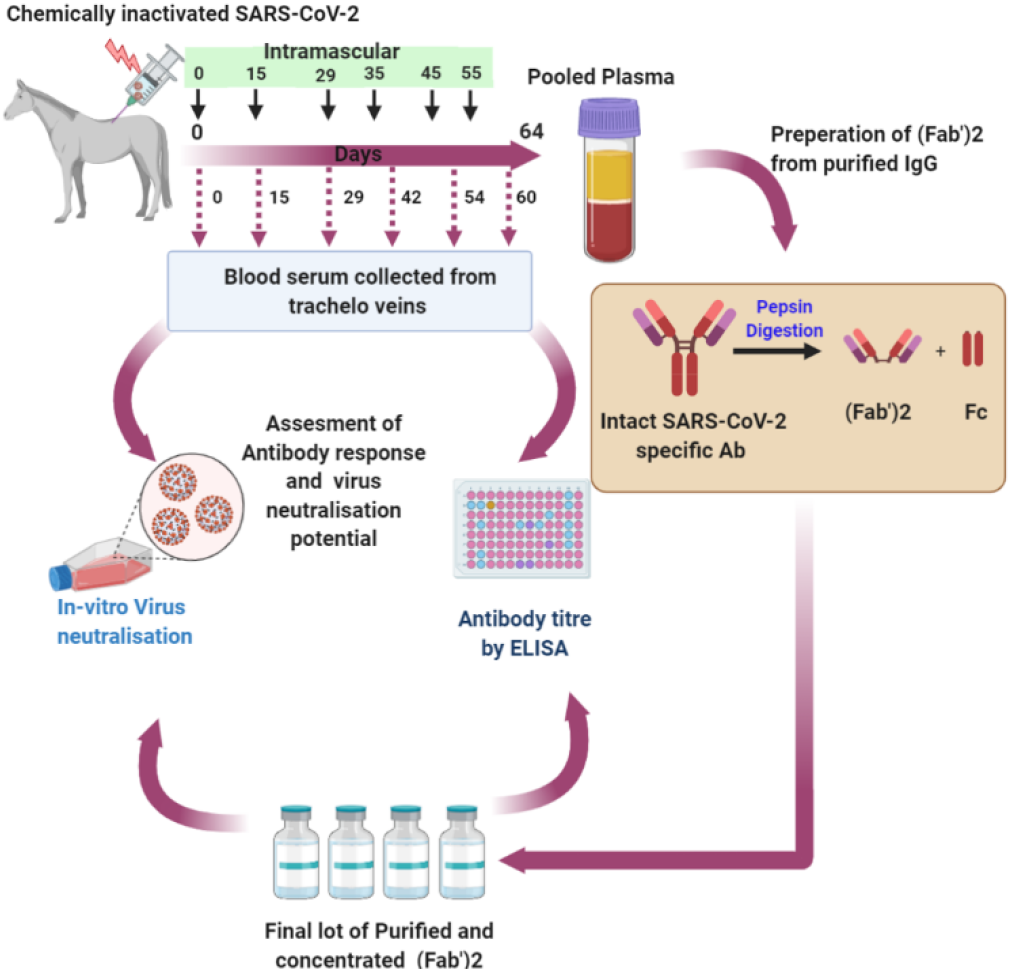
Immunization scheme and workflow. BPL-inactivated SARS-CoV-2 particles were mixed with FCA and injected intramuscularly into the equines. Immunization was repeated on the days mentioned in the scheme. Plasma collected from the immunized animals were pooled, their antibody response was assayed and virus neutralization titer was quantified by microneutralization assay. Subsequently, IgG was purified from the pooled plasma, digested with pepsin and the F(ab’)_2_ fragment was purified. Neutralization titers of these purified and concentrated fragments were also assayed.

## RESULTS

### Isolation of SARS-CoV-2 particles and establishment of virus culture

Out of several cultures established, two cultures were used for this study. The cultures continued to demonstrate the presence of the virus in the supernatant as suggested by qRT-PCR (data not shown). Plaque forming assay revealed high titers of infectious virus particles in the order of 10^7^ PFU/mL in these supernatants (data not shown). This culture was further expanded to larger size and stored for subsequent studies. To verify the presence of SARS-CoV-2, we analyzed the presence of virion proteins in the supernatants. Supernatants containing infectious viral particles were precipitated by ultracentrifugation, lysed, and subsequently subjected to immunoblotting. As demonstrated in Figure 2A, the spike (S) and nucleocapsid (N) proteins of SARS-CoV-2 were detected in the concentrated viral supernatants confirming the presence of the virus. In parallel, immunoblot analysis of SARS-CoV-2 infected Vero cells detected the robust expression of S and N proteins, further confirming the establishment of active SARS-CoV-2 cultures (Figure 2B). The viral genome sequences are available at GISAID (hCoV-19/India/TG-CCMB-O2-P1/2020 [hereafter referred to as CCMB-O2], GISAID accession-EPI_ISL_458075) and hCoV-19/India/TG-CCMB-L1021/2020 [referred to as CCMB-L-1021], GISAID accession-EPI_ISL_458046). We then examined the inactivation of the virus by β-propiolactone (BPL). We used either 1:250 or 1:1000 dilutions of BPL (v/v in media) in this study, both of which displayed total inactivation of the virus. Detection of viral proteins S and N confirmed the retention of the protein integrity of the inactivated viral stocks to induce immune response in the equines (Figure 2C).

**Fig.2.**
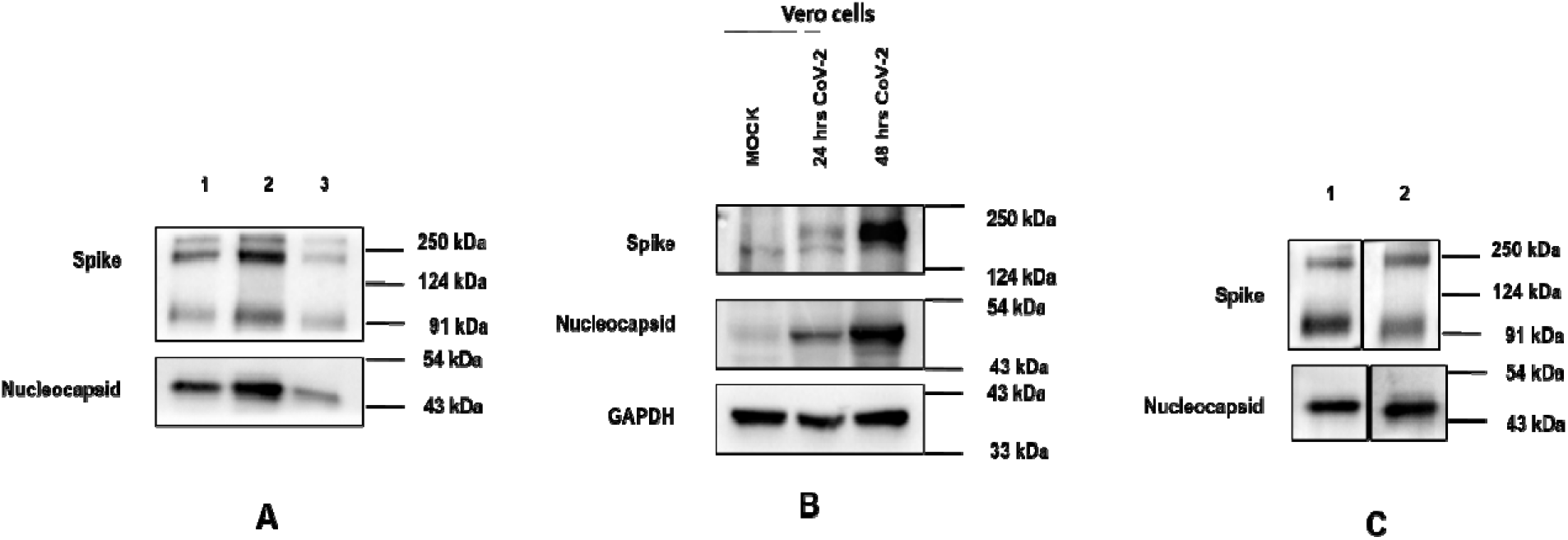
Characterization of SARS-CoV-2 isolate supernatant and *in vitro* infection. (A) Immunoblotting of SARS-CoV-2 spike and nucleocapsid proteins in three independent supernatants from *in vitro* cultures of Vero cells. Briefly, Vero cells were infected with SARS-CoV-2 stocks at 1:10 dilution and three days later supernatants were collected. 10 mL of three independent supernatants were ultra-centrifuged at 100000 × g for 90 minutes and the pellets were re-suspended in 1 mL of PBS, lysed with 2 × lysis buffer, and immunoblotted. Results from three independent supernatants are depicted. (B) Expression of spike and nucleocapsid proteins in Vero cells infected with SARS-CoV-2. The cells were harvested either at 24 or 48 hrs post-infection before subjecting to immunoblotting. (C) Detection of spike and nucleocapsid in BPL-treated viral supernatants. The supernatants were precipitated as in (2A) after the inactivation with BPL. Two individual samples were processed for immunoblotting.

### Antigen-specific immune response

As explained in the Methods section, the horses were injected with inactivated SARS-CoV-2 and blood samples were collected periodically. Plasma/sera prepared from individual animals were subjected to ELISA to quantify IgG levels. Inactivated viral antigens induced strong IgG response from 29^th^ day onwards peaking at 42 days after priming and subsequently stabilizing as shown in the Figure 3A. Notably, 80% of the immunized horses showed the seroconversion from 29^th^ day onwards except two animals which remained non-responsive during the entire period of study (Figure 3B). The antibody titer which is indicative of quality of induced antibody also enhanced from 29^th^ day (1: 25600 dilution) as compared to the negative control, peaking at 42^nd^ day post immunization (1:51200) and later retreating to 1: 25600 at 54^th^ day as demonstrated in the Figures 4 A and B.

**Fig.3.**
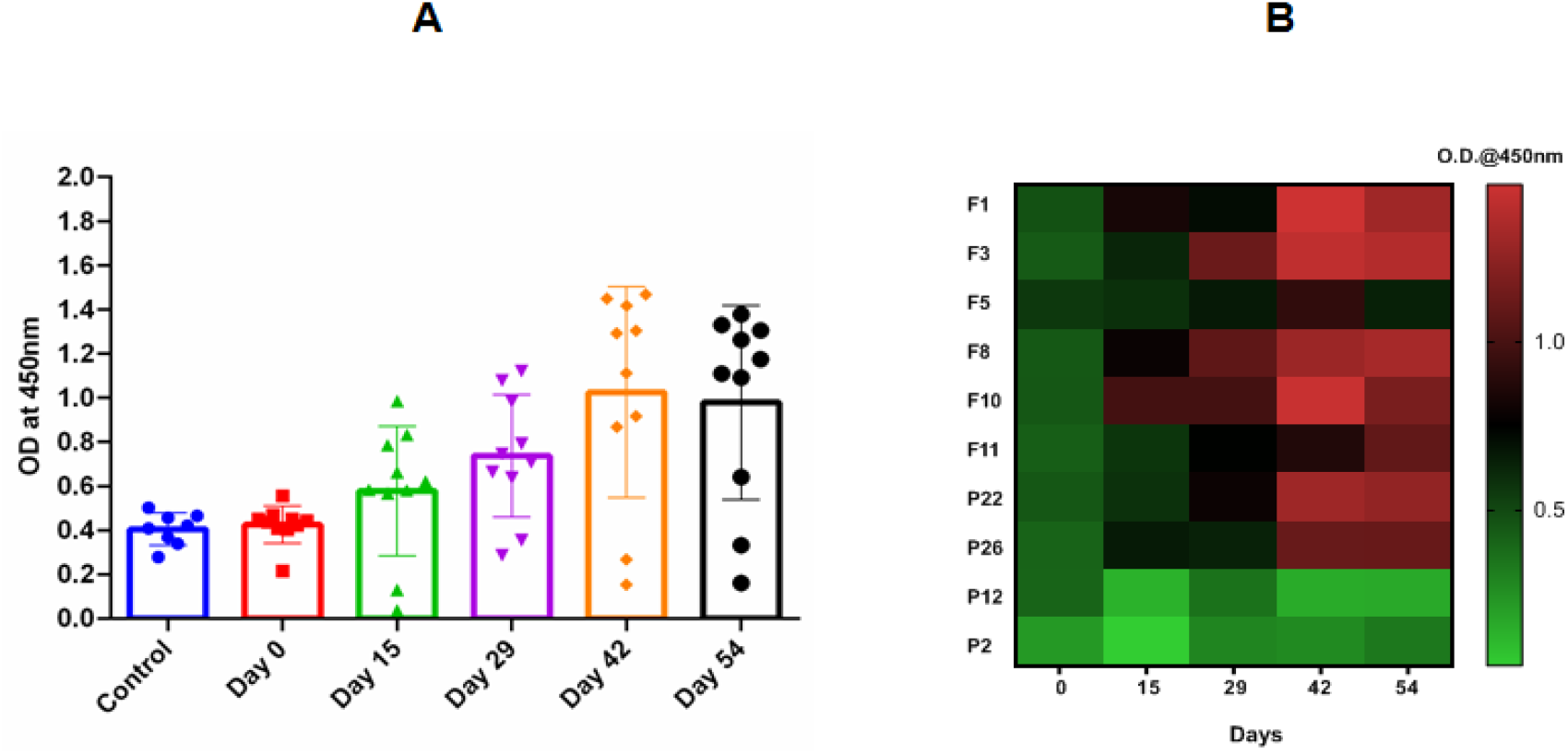
Evaluation of SARS-CoV-2 specific total IgG from serum collected at specified time points after first immunization using indirect ELISA. A) Antigen response kinetics of 10 individual horses along the course of time (Day 0 to day 54) with respect to control (pre-immunized sera). B) Heat map of the same with labeled individual animal.

**Fig.4.**
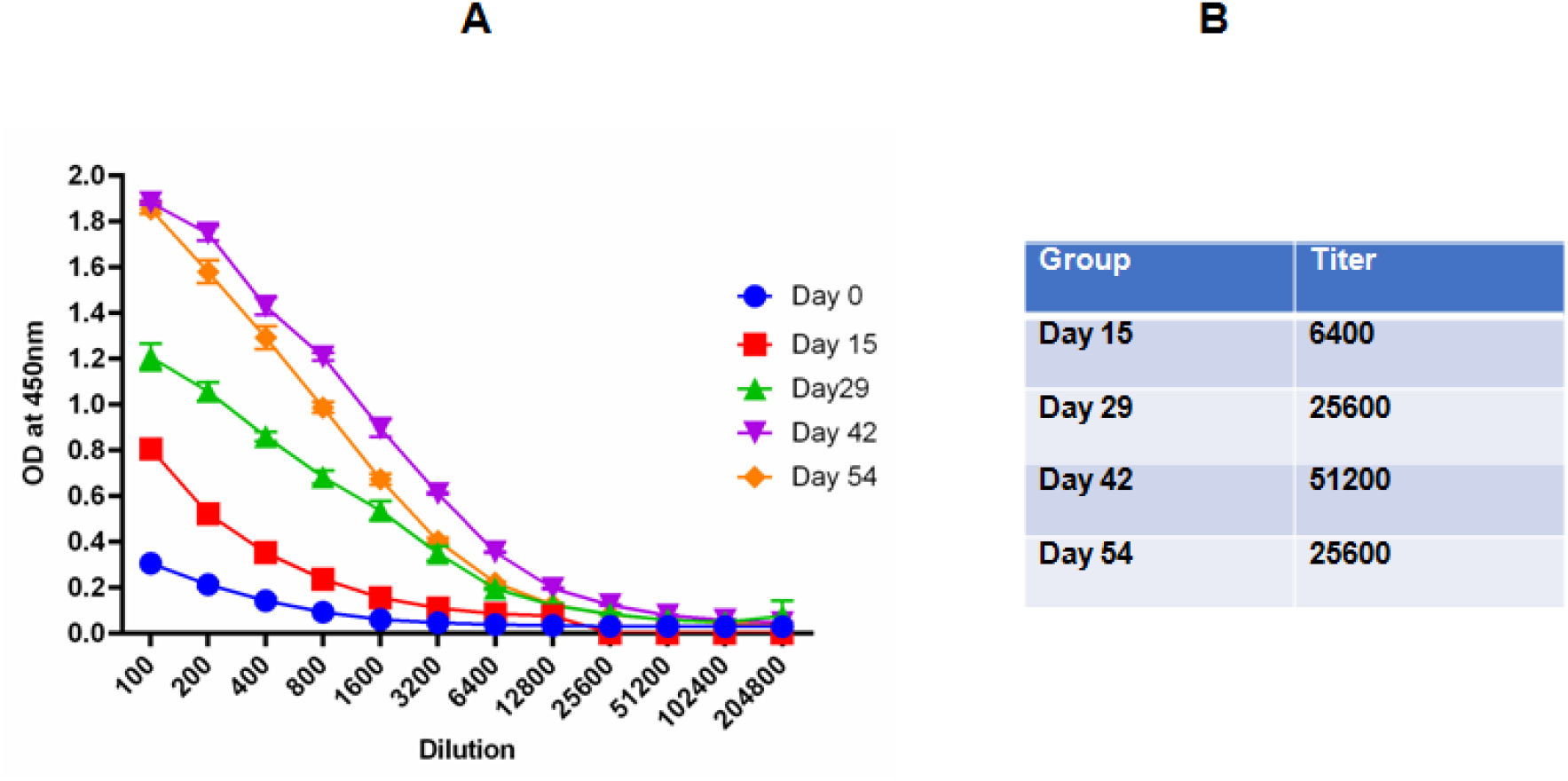
Antibody titration kinetics of serum collected along the different time points using indirect ELISA. A) Serially diluted serum (100 to 204800) used over the virus antigen coated ELISA plate and absorbance value at each dilution and time points represented at Y axis. B) Antibody titers were calculated by the reciprocal value of highest dilution at which absorbance value is ≥ twice the value of negative control in the same dilution series.

### Characterization of F(ab’)_2_ and measurement of their binding titer

Pepsin treatment of the purified IgG-generated F(ab’)_2_ fragments and the purified F(ab’)_2_ fraction showed the characteristic peak in the chromatogram (Figure 5A) with a typical band visible around 110 kDa region in the non-reducing condition and 25 kDa in the reducing condition (Figure 5B), demonstrating the purity of F(ab’)_2_ preparation. In the non-reducing and reducing condition, F(ab’)_2_ typically shows single band at ∼110kDa and 25 kDa position respectively whereas whole immunoglobulin shows a single band at 150 kDa in non-reducing condition and two bands at 75 kDa (heavy chain) and 25 kDa (light chain) positions under reducing condition. This result confirms that immunoglobulin has been successfully converted into F(ab’)_2_ fragments. Next, we measured the titer of the purified F(ab’)_2_. The purified F(ab’)_2_ samples showed a remarkable titer of 1:102400 as compared to the negative control (Figure 6).

**Fig.5.**
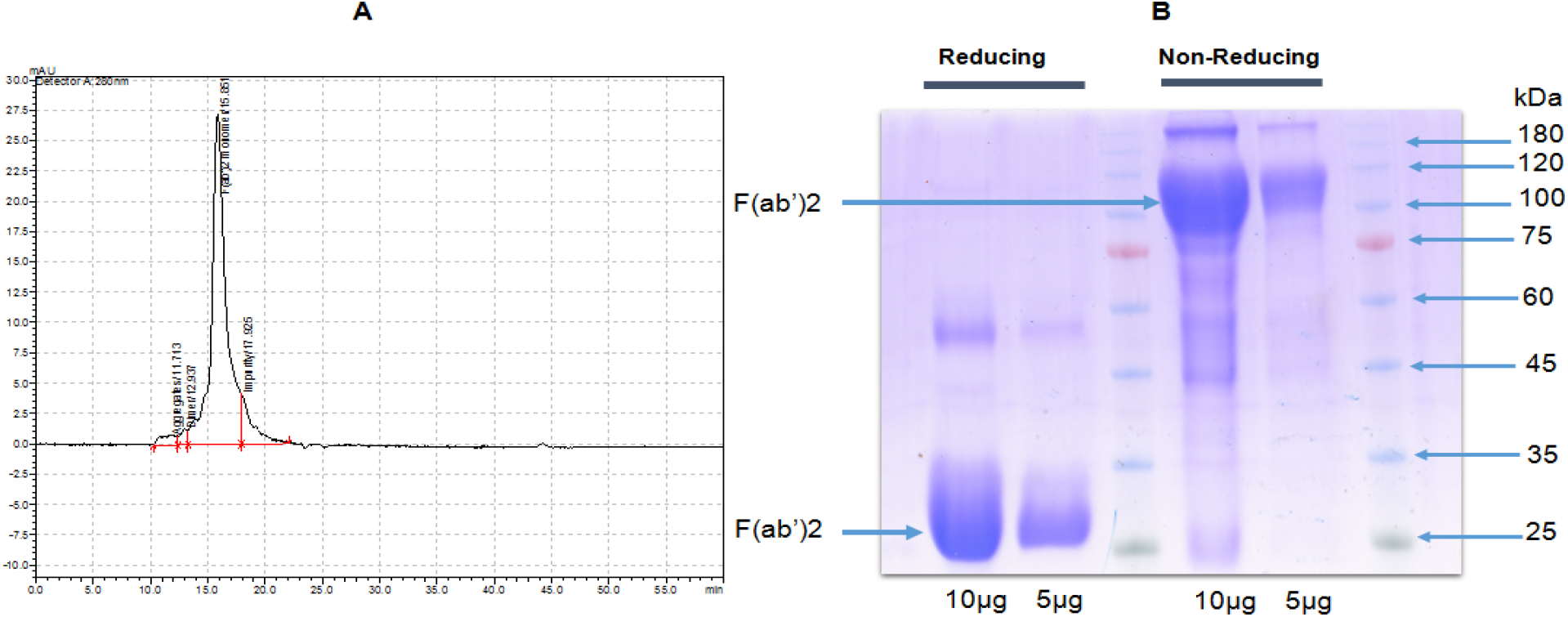
Characterization of purified F(ab’)_2_. A) HPLC chromatogram shows a dominant F(ab’)_2_ peak B) SDS-PAGE image of purified F(ab’)_2_ under reducing and non-reducing conditions. The purified F(ab’)_2_ were loaded (5 or 10 µg) with and without beta-mercaptoethanol in SDS-polyacrylamide gel and resolved under constant voltage. Reducing and non-reducing conditions were achieved with and without beta-mercaptoethanol respectively. The result shows F(ab’)_2_ fragment of 28 kDa (two heavy chains and two light chains of almost similar molecular weight) under reducing condition (left) and >110 kDa under non-reducing condition (right) demonstrating the purity of F(ab’)_2_ preparation.

**Fig.6.**
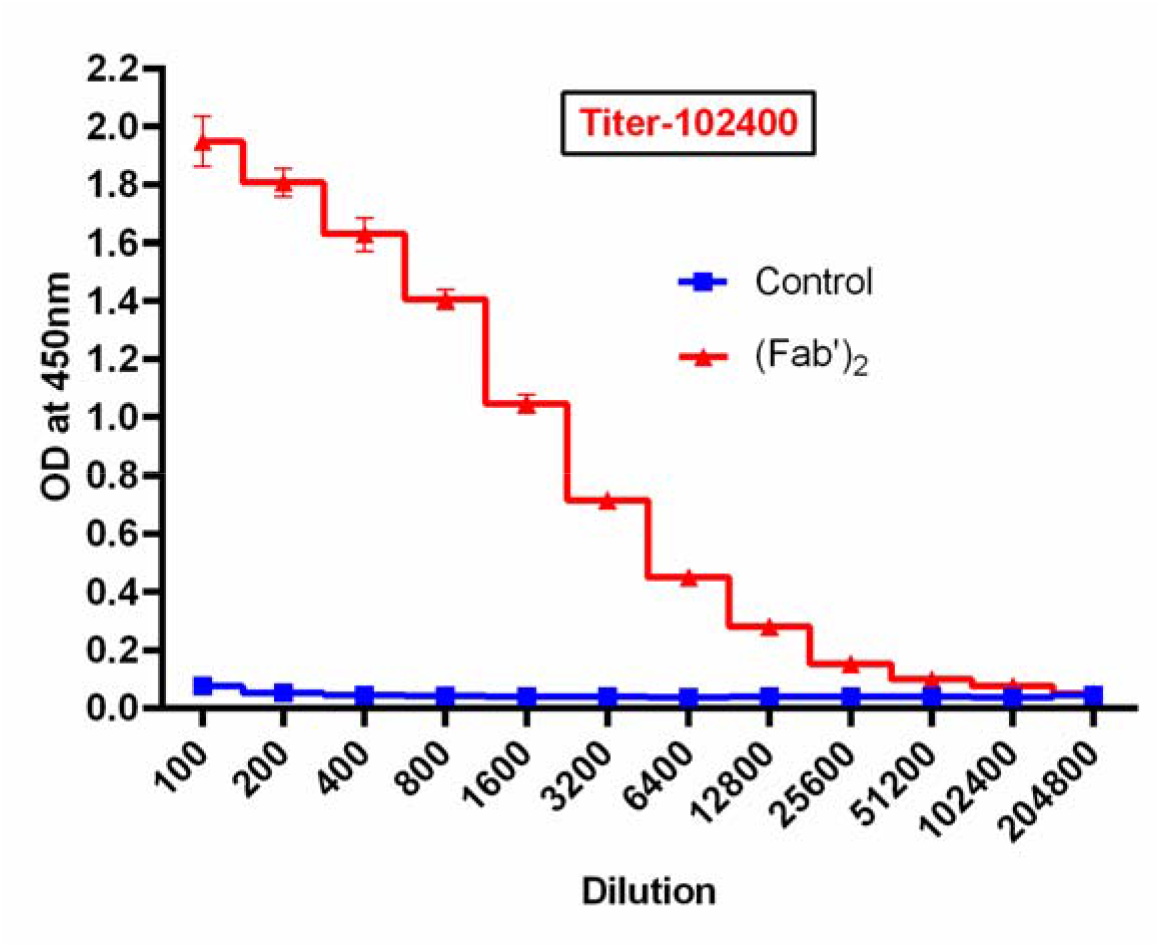
Antibody titration of the F(ab’)_2_ purified from the pooled plasma of immunized equine with respect to pooled sample of negative control and titer is given in inset box. F(ab’)_2_ titer was measured by direct ELISA method in which the whole virus antigen (approximately 1×10^5^ virus particles) coated-plates were incubated with serially diluted F(ab’)_2_ (1:100 to 1:204800 dilution) for 2 hours at RT. HRP conjugated anti-F(ab’)_2_ secondary antibody (1:5000 dilution) were added subsequently and the color reaction was developed by adding TMB substrate. F(ab’)_2_ titer was calculated by the reciprocal value of highest dilution at which absorbance value is ≥ twice the value of negative control in the same dilution series.

### **Neutralization of SARS-CoV-2 by antisera and purified** F(ab’)_2_

Antisera from five individual animals were pooled and the virus neutralization capacity was quantified by microneutralization assay. Sera from days 29, 42 and 54 were identified for neutralization assay. The pooled sera were serially diluted at 1:2 ratio in serum-free media and each dilution sample was incubated with Vero cells for infection. As demonstrated in Figure 7A, the antisera from three independent time points displayed significantly higher neutralization capacity over the control sera. Significant neutralization capacity was achieved 29 days post-immunization and spiked at 42 days post-immunization (Figure 7A).

**Fig.7.**
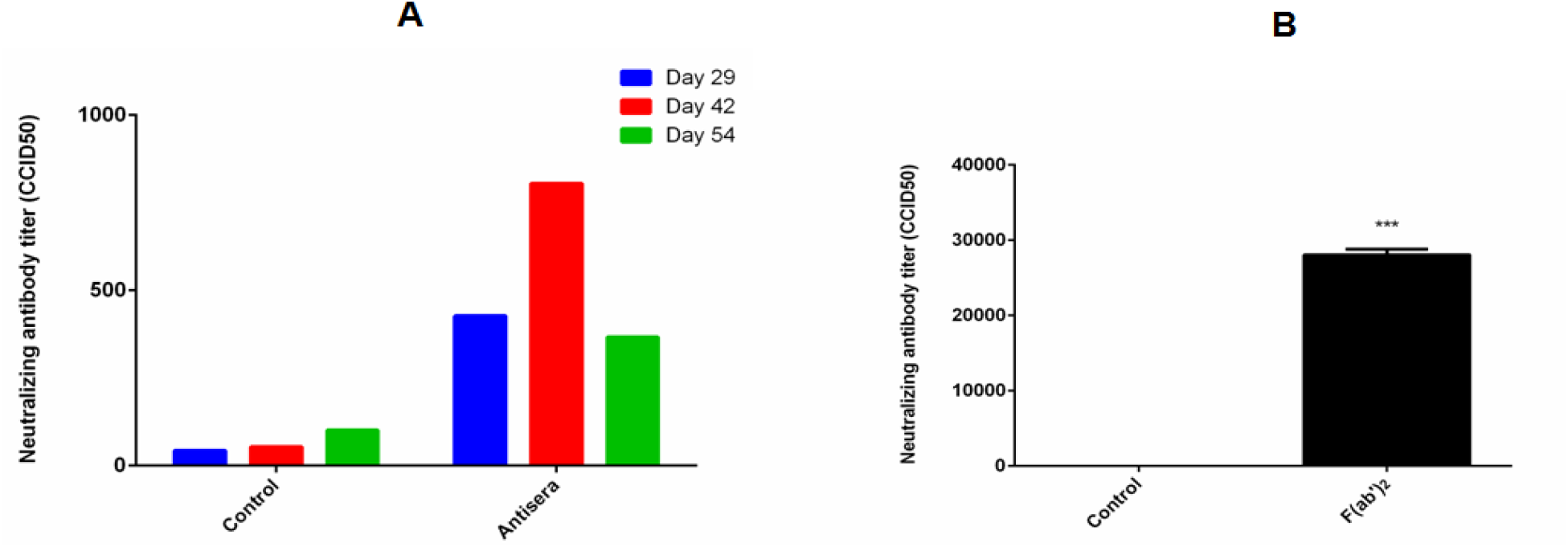
Neutralization capacities of host antisera and purified F(ab’)_2_. (A) Neutralization of SARS-CoV-2 by pooled of antisera. Neutralization capacities of antisera drawn from horses 29, 42 and 54 days post-immunization were tested by micro neutralization assays. CCID50 of the antisera treated virus particles are represented. (B) Neutralization capacities of F(ab’)_2_ generated from pooled antisera. Micro neutralization assays were performed similarly as in Figure 6 and the data are represented as CCID50

Next, we assayed the neutralization capacity of purified F(ab’)_2_ fragments from the antisera samples. Here, three separate F(ab’)_2_ pooled batches of antisera were assayed for neutralization. As demonstrated in Figure 7B the purified F(ab’)_2_ achieved significantly high neutralization titers well above 25000. We also tested the neutralization capacity against another strain CCMB-L-1021 that contained D614G mutation in its Spike protein. The antisera generated against CCMB-O2 demonstrated reasonably high neutralization titer against CCMB-L-1021 containing D614G, albeit lower than that against CCMB-O2, indicating the efficacy of polyclonal antisera against other variant strains of SARS-CoV-2 (Supplementary Figure 1). Cross neutralizing ability of antisera reduces the risk burden of its therapeutic relevance against emerging SARS-CoV-2 variants, thereby suggesting that the equines purified F(ab’)_2_ based passive immunotherapy hold enormous therapeutic potential against COVID19 in terms of cost, safety, storage and mass availability.

## DISCUSSION

Emerging and re-emerging zoonotic viral infectious diseases such as SARS-CoV-2, SARS-CoV and MERS-CoV have become more frequent in the recent past due to the ever-increasing encounters with wild animals and pose great threat to public health system. Despite great advancement in the field of biotechnology and pharmaceutics, effective response against such kind of pandemic is still lagging. Several vaccines have been at the threshold of being introduced into the markets and they are reported to be quite effective while some of them have been approved for emergency use ^3,24^. Nevertheless, antibody therapy still holds important position in the fight against COVID-19 since vaccinating the entire human population would require years of continuous vaccination. Several monoclonal antibodies (mAbs) have shown their neutralization potential against SARS-CoV-2 ^25,26^, but production of individual mAbs are resource-exhaustive and also bear the risk of losing their potential against the possible mutants in the specific epitopes. Polyclonal antibodies generated in large animals such as equines have the advantage of faster generation, requirement of relatively much smaller investment and efficacy against multiple epitopes. Antisera generated from such sources had been a great source of antiviral antibody to treat the various viral infection such as SARS-CoV, Ebola, MERS-CoV and avian influenza virus^27–30^. Clinical evidence of COVID-19 disease shows that latent period of infection is short and majority of the patients recover faster without any persistent infection thus increasing the prospects of using neutralizing antibodies in blocking the SARS-CoV-2 virus particles^31^. Even though convalescent plasma from the recovered patients was considered to be a great source of neutralizing antibodies^32^, the difficulty in recruiting such individuals along with the lack of consistency in the neutralizing antibody titer among them has posed major obstacles in utilizing its potential^33^. Moreover, several reports indicate the lack of efficacy of convalescent plasma therapy in improving the severity of COVID-19 ^34,35^.

Notwithstanding the use of plasma therapy to treat various infectious viral disease such as SARS, H5N1 and Ebola,^28,36,37^ it always carries a risk of blood-borne infection and its limited availability hampers its prospect of universal application.

Considering the enormous potential of antibody based therapy, we developed a SARS-CoV-2 specific immunoglobulin fragments F(ab’)_2_ in equines using chemically inactivated virus as similar to other reported work^12,28^. In the current study, we report the serum IgG titer to 1:51200 at 42 days’ post immunization, which is significant as compared to earlier report against the SARS-CoV^28^. Another study demonstrated the potency of virus like particle (VLP) of MERS-CoV in the equine ^27^ where they attained the antibody titer of 1: 20480. The higher antibody response in this study might be due to the optimum antigen dose and immunization schedule.

The immunoglobulin fragment is normally developed by the proteolytic cleavage of immunoglobulin that potentially negates the side effect of serum sickness and antibody dependent enhancement of infection (ADE) as it no more binds to the Fc receptor of the immune cells. Hence, it is suitable for usage at larger doses without any off-target concerns^38^. F(ab’)_2_ that we generated achieved greater antigen specific antibody titer of 1: 102400 which is comparatively better the earlier published reports ^27,28^. Other recent reports also demonstrated high antibody titer for F(ab’)_2_ generated from horses using receptor binding domain (RBD) of spike protein^18,19^, but *in vivo* response still needs to examined before any conclusion. Generation of inactivated virus was more direct and feasible approach for us than raising large amounts of vaccine quality spike proteins to be used at commercial levels. Apart from the lesser side effect, F(ab’)_2_ can penetrate deeper into the organs due to smaller size and lesser cellular affinity therefore it can neutralize the virus in the extravascular tissue ^39^.

Virus neutralization by an antibody is the major goal of antiviral passive immunotherapy as the effective antibody should neutralize the virus *in vitro* and *in vivo* setup at a significantly higher dilution. At this front, F(ab’)_2_ generated in this study demonstrates robust *in vitro* virus neutralization titer as high as 28000 which is comparable with other similar studies against the SARS-CoV-2^18,19,28^. Effective neutralization of a variant carrying mutation in spike protein by this antisera demonstrates the broad efficacy in using purified antisera of equine origin. Furthermore, the F(ab’)_2_ antibody shows very significant virus neutralization that is comparable or higher than that the convalescent plasma therapy offers without the risk of blood born disease^40^. The F(ab’)_2_ antibody developed from the whole virus antigen contains polyclonal antibody against all possible antigens exposed, having a broader range of binding repertoire thereby providing better anticipated scope of virus neutralization as compared to subunit polyclonal antibody or monoclonal antibody.

The current study is based on the tested method for production of antigen specific antibody in equines and is easy to scale up by industry with the domain expertise. The WHO guidelines are already laid out for the production and application of antisera and their product from equine source therefore it can be quickly available to the world for immediate application^41^. Strain-specific antisera can be developed quickly based on the necessity. Hence F(ab’)_2_ antibody from hyper-immunized equine serum could be a viable option for passive immunotherapy to treat the COVID-19 and along with vaccines, can collectively bring down the burden of the pandemic. Even in the optimistic scenario of active vaccines, passive immunotherapy can also be used to save the life of terminally ill patients as it has been used to treat rabies.

## Conclusion

Anti-SARS-CoV-2 immunoglobulin F(ab’)_2_ prepared from the plasma of hyper-immunized equines demonstrated high antibody titers and effective neutralization of the parental as well emerging viral strains would be a reliable and efficient tool in the fight against COVID-19.

## Author Contributions

D.G. optimized large-scale SARS-CoV-2 virus propagation, BPL inactivation and microneutralization assay. D.G., and D.K propagated, quantified and inactivated large-scale SARS-CoV-2 cultures, and performed microneutralization assays. D.V. and V.S established SARS-CoV-2 cultures used in this study. H.P. performed immunoblotting. N.K. conceived and conceptualized the study along with K.H.H. F.A. and R.K. performed the immunological characterization of anti-sera and F(ab’)_2_. D. G., F.A., K.H.H. and N.K. wrote the manuscript. C.K., P.S., and S.K performed the equine immunization and F(ab’)_2_ preparation. S.R provided the patient sample for the isolation of SARS-CoV-2.

## Acknowledgement

Several volunteers at the Centre for Cellular and Molecular Biology who were part of COVID-19 testing are thanked for their help in identifying SARS-CoV-2 positive samples for virus culturing. Mohan Singh Moodu and Amit Kumar contributed significantly with the logistics. We thank Karthika Nair, Abhirami P S, and Sai Poojitha for their help with various experiments.

## Institutional ethics clearance

Institutional ethics clearance (IEC-82/2020) was obtained for the patient sample processing for virus culture.

## Institutional biosafety

Institutional biosafety clearance was obtained for the experiments pertaining to SARS-CoV-2.

## Supplementary Figure 1

**Fig.S1.**
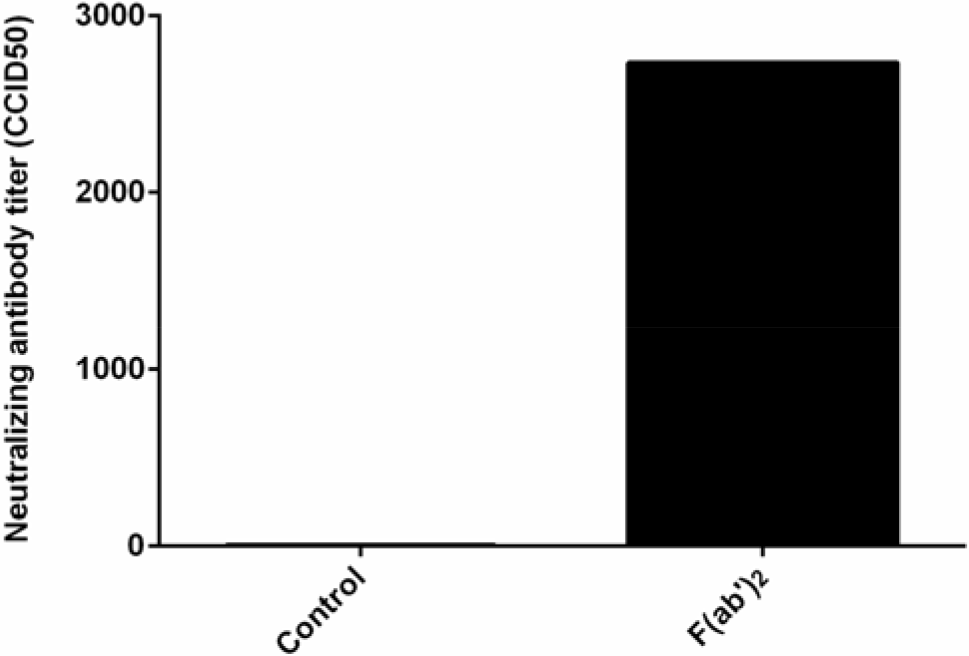
Neutralization capacities of F(ab’)_2_ generated from batch of pooled antisera. Micro neutralization assays were performed similarly as in Figure 6 and the data are represented as CCID50. The antisera were developed against CCMB-O2 isolate and neutralization was tested against CCMB-L-1021.

